# Engineered Matrices Reveal Sulfation-Mediated Stress Adaptation and Drug-Specific Modulation of Chemotherapeutic Response

**DOI:** 10.64898/2026.05.21.726894

**Authors:** Sevgi Sarica, Ece Öztürk

## Abstract

Engineering biomimetic extracellular matrices that isolate specific biochemical cues is essential for understanding how matrix chemistry regulates tumor cell behavior and therapeutic response. Aberrant sulfation due to proteoglycan expression is a hallmark of lung tumor matrices, yet its functional impact is difficult to study using conventional materials where mechanical and biochemical variables are coupled. To address this, mechanically matched sulfated alginate hydrogels are engineered to mimic the elevated sulfated glycosaminoglycan (sGAG) content of malignant ECM, enabling sulfation to be examined as a single, tunable variable. Within this system, ECM sulfation is shown to enhance tumor cell proliferation, promote oxidative and mitochondrial stress tolerance, suppress apoptotic signaling and attenuate the efficacy of cisplatin, gemcitabine and paclitaxel. Sulfated matrices preserve mitochondrial membrane potential, limit ROS accumulation, shift apoptotic gene expression toward a survival-favoring profile, selectively upregulate ABCB1-mediated efflux and modulate drug response through the PI3K/Akt-ABCB1 signaling axis. Functional inhibition of PI3K and ABCB1 uncovers drug-specific dependencies while dual pathway targeting completely restores chemotherapeutic sensitivity. These findings identify ECM sulfation as a potent regulator of stress adaptation and therapeutic efficacy in lung adenocarcinoma and underscore the importance of biomimetic ECM design in controlling tumor cell fate and drug response.

## 1. Introduction

Lung cancer is the leading cause of cancer-related mortality worldwide [1]. Non-small cell lung cancer (NSCLC) is the most prevalent form with almost 85% incidence among lung tumors and a poor 5-year survival rate [2]. Albeit advancements in therapeutic regimes, overt clinical symptoms occur at later stages and cause low response to therapy [3]. Surgical resection remains a successful option for early-stage cases, while administration of platinum-based combinational chemotherapy and targeted therapy for locally advanced or metastatic lung cancers has poor clinical outcome due to low response and resistance [3–6]. Hindrance in therapeutic efficacy may be attributed to several mechanisms including increased drug efflux, cell adhesion-mediated suppression of apoptosis, alteration of drug metabolism, cancer stemness, enhanced DNA repair, activation of alternative signaling pathways and microenvironmentally induced dormant state [7–13]. Controlling the abovementioned molecular barriers calls for a comprehensive understanding of the interplay between tumor cells and their surrounding microenvironment in order to unveil novel strategies to improve therapeutic response, overcome resistance development and increase survival rates [14].

The tumor microenvironment (TME) is a complex niche which acts as a determinant for tumor growth and dynamically evolves throughout disease progression as well as treatment leading to heterogeneity in therapeutic response among patients [15–18]. Studies on mechanisms of drug resistance in cancer have long relied on conventional monolayer culturing of tumor cells which lacks a representation of the complexity of the TME and favors rapidly proliferating populations leading to overestimation of therapeutic efficacy [19–21] Genetically engineered mice and patient-derived xenografts (PDX) provide system-level models, however, offer limited genetic and environmental manipulation together with high costs and low throughput [18, 22]. The extracellular matrix (ECM), a key component of the TME, is an important regulator of tumor cell interactions with other cell populations including fibroblasts, endothelial cells and immune cells [23, 24]. The need for faithfully recapitulating the tumor ECM characteristics with controlled tunability and enabling cell-matrix interactions within human tumor models led to the convergence of cancer research and tissue engineering [25]. Engineered 3D tumor models utilizing naturally derived scaffolds such as collagen or reconstituted basement membrane (rBM) and synthetic materials such as polyethylene glycol have been developed [26–28]. These models allowed isolation of specific aspects of the intricate tumor ECM network and revealed their effect on tumorigenesis, invasion and responses to chemotherapy [29–31]. The importance of matrix stiffness and cell adhesive ligand content in regulation of malignant progress and therapeutic responses have been established with seminal studies demonstrating the need for representative 3D human tumor models to perform clinically relevant drug screens [32–35]. Patient-derived organoid cultures in rBM have emerged with ability to maintain cellular heterogeneity as well as morphological features and genetic background of native tumor revolutionizing drug screening studies in lung cancer [36–38].

Lung tumors exhibit distinct changes in ECM composition compared to healthy lung tissue marked with an increase in cell instructive ECM ligands [39]. Among those ligands, proteoglycans (PGs) hold a vital role in regulating tumor growth and progression through enabling receptor tyrosine kinase (RTK) signaling [40–46]. PG structure consists of a core protein with covalently linked brush-like sulfated glycosaminoglycan (sGAGs) side chains [47]. The negatively charged sulfate groups on sGAGs confer affinity to growth factors and cytokines while serving as their coreceptors via mediating their binding to RTKs [45]. Our group recently showed that PG genes are aberrantly expressed in NSCLC patient tumors correlating with increased invasive potential, expression of stemness markers and poorer survival [43]. Similarly, suppression of a sulfate transporter was shown to re-sensitize resistant melanoma with high YAP1 activity against MAPK inhibitors [48]. Transmembrane PG syndecan-1 has been reported to induce stemness in inflammatory breast tumors leading to chemoresistance [49]. Expression of NG2, a transmembrane chondroitin sulfate PG, has been shown to correlate with progenitor population and promote chemoresistance via the integrin-dependent PI3K/Akt signaling axis in brain tumors [50]. Yet, the role of extracellular PG/sGAG enrichment in the TME on chemoresistance is still largely unknown.

In present work, we investigated the role of PG/sGAG enrichment in tumor ECM on chemotherapeutic response of lung cancer cells within a biomimetic 3D human lung tumor model. Sulfation in the lung TME leads to a significant reduction in the response of tumor cells to clinically used therapeutics cisplatin, gemcitabine and paclitaxel. Tumor cells in sulfated hydrogels demonstrated significant suppression of apoptosis genes as well as upregulation of genes regulating survival and drug export. Loss-of-function assessment revealed different modes-of-action for each drug and distinct routes of re-sensitization of tumor cells. Our work exhibits that increased sulfation acts as a crucial ECM parameter that diminishes response to chemotherapeutics in the lung TME in a drug-specific manner. The findings highlight the importance of developing engineered tumor models for elucidating the biological mechanisms of ECM-induced resistance to therapeutics and forming strategies to re-sensitize tumor cells.

## 2. Results

### 2.1. Sulfation of Alginate Promotes Enhanced Proliferation of Lung Adenocarcinoma Cells

In order to model the increase of PGs/sGAGs in the lung TME, we pursued chemical modification of alginate with sulfate moieties. We have previously shown that alginate, an inert and biocompatible polymer which lacks cell adhesion cues, can successfully mimic native sGAGs through its sulfation which enables growth factor entrapment and mediation of ligand interactions with receptor tyrosine kinases (RTKs) [42, 43]. This strategy allows for both tunability in the degree of sulfation and independent control over hydrogel stiffness [42, 51]. To study the sole effect of extracellular sulfation on chemotherapeutic response of lung tumor cells, we constructed either unmodified alginate (Alg) hydrogels representing an inert microenvironment as well as tumor-mimetic sulfated alginate (S-Alg) hydrogels recapitulating the abundant presence of proteoglycans in tumor matrices (**Figure 1A**). Hydrogel stiffness was tuned with the ionic crosslinker (Ca^2+^) to match the lung tissue microenvironment as the tumor’s site of origin [42, 43]. Oscillatory rheology measurements were performed to assess storage modulus (G’) and loss tangent (G’’/G’) in both hydrogels and did not show a significant difference in stiffness and gelation capacity in Alg and S-Alg matrices (**Figure 1B**).

**Figure 1.**
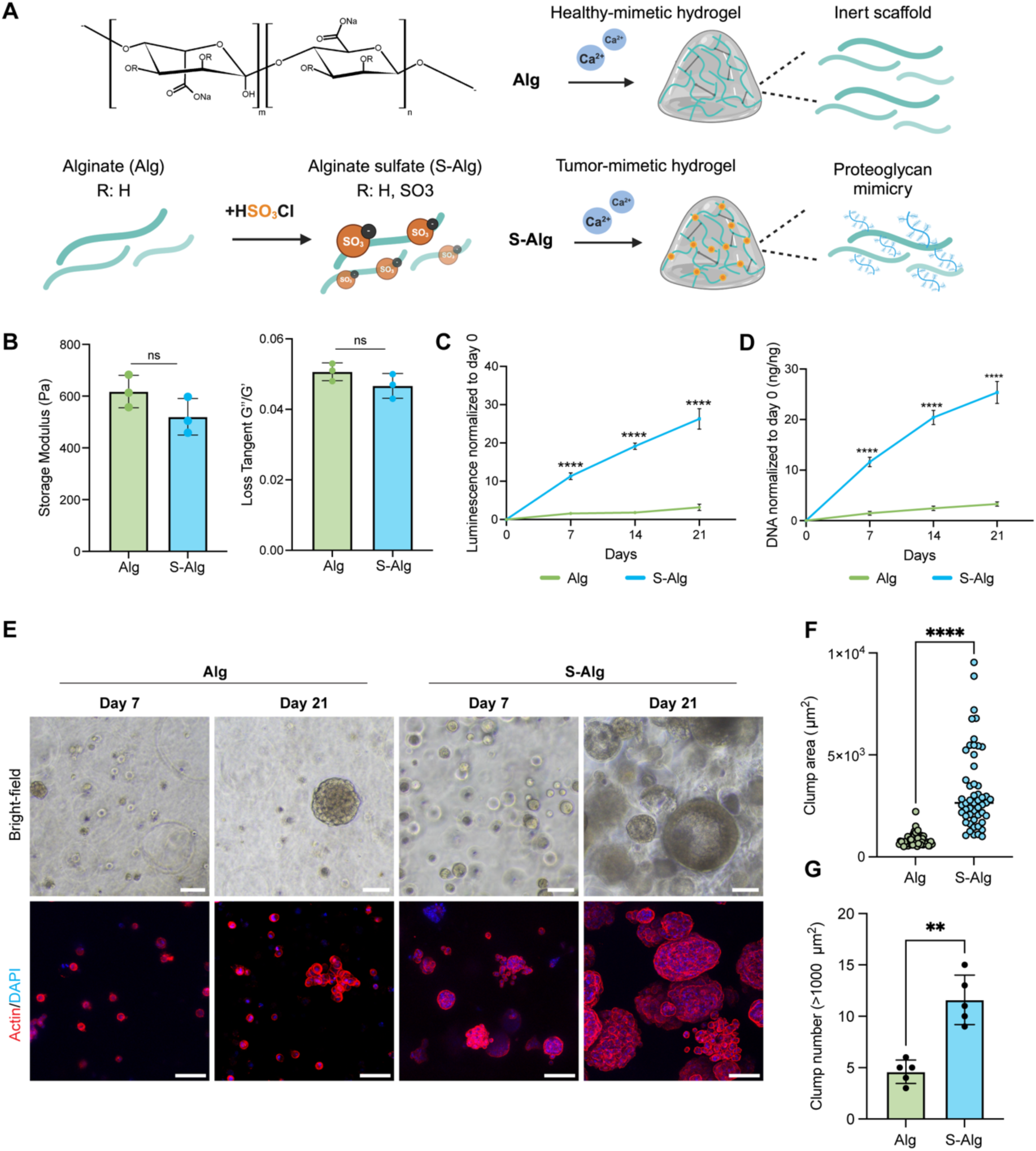
Mimicking sGAG enrichment in the extracellular microenvironment promotes proliferation of lung adenocarcinoma (A549) cells. **A)** Schematic of alginate modification and fabrication of alginate (Alg) and sulfated alginate (S-Alg) hydrogels with Ca^2+^ crosslinking. **B)** Storage modulus (G’) and loss tangent (G’’/G’) of Alg and S-Alg hydrogels assessed with oscillatory rheology measurements. Three independent replicates were measured with error bars indicating mean ± SD. Statistical analysis was performed using two-tailed t-test, ns not significant. **C)** Metabolic activity and **D)** dsDNA quantification of A549 cells encapsulated in Alg and S-Alg hydrogels at days 0, 7,14 and 21. Three independent replicates were measured, and error bars indicate mean ± SD. Statistical analysis was performed using two-tailed t-test, ****p<0.0001. **E)** Bright-field (upper panel) and phalloidin (red)/DAPI (blue) staining (lower panel) images of A549 cells encapsulated in Alg/S-Alg hydrogels at days 7 and day 21. Scale bar:100 µm. **F)** Quantification of clump area and **G)** clump number (>1000 µm^2^) of cells grown in Alg and S-Alg hydrogels at day 21. Data are shown as mean ± SD of five replicates (the average of 10-30 images per replicate). Statistical analysis was performed using two-tailed t-test, ****p<0.0001, **p<0.01.

Next, non-small cell lung adenocarcinoma cells (A549) were encapsulated in Alg and S-Alg hydrogels at a density of 2x10^5^ cells/ml and cultured for 3 weeks in order to monitor the effect of sulfation on cell growth and metabolic activity (**Figure 1C**). Sulfation had a potent effect on lung tumor cell growth and tumor clump formation with a significant increase in metabolic activity in S-Alg hydrogels at all time points (**Figure 1C**). dsDNA content quantification was performed to validate cellular proliferation and similarly revealed a significant increase in S-Alg hydrogels (**Figure 1D**). Clump formation of tumor cells encapsulated in Alg and S-Alg hydrogels was monitored at days 7 and 21. Bright-field images and phalloidin/DAPI stainings of hydrogels revealed distinct changes in cell density, clump size and morphology in S-Alg matrices (**Figure 1E**). Analysis of cellular clusters grown in S-Alg matrices showed significantly higher clump size as well as clump number (**Figure 1F, G**). Sulfation supported formation of larger and more cohesive clumps with continuous cell-cell contacts and a smooth peripheral contour whereas single cell growth was dominantly observed in inert Alg hydrogels. Our results demonstrated that sulfation alone is a strong inducer of tumor cell growth in an otherwise inert microenvironment.

### 2.2. Growth in 3D Sulfated Environment Potently Reduces Tumor Cell Sensitivity to Chemotherapeutics

We next sought to understand the role of aberrant sulfation in the cellular microenvironment on tumor cell response to chemotherapeutics. Cisplatin, gemcitabine and paclitaxel were used in the study as they all are frequently used in the clinics for lung cancer treatment [2, 4]. Initially, we performed a dose-response screen for all drugs in conventional monolayer culture of lung tumor cells (**Figure 2A**). A549 cells were seeded onto 96-well plates with a density of 3x10^3^ cells/well and cultured overnight prior to treatment with the chemotherapeutics at indicated doses. In line with literature, half-inhibitory doses (IC_50_ values) of all drugs for 2D culture were determined as 10.16 µM, 4.7 µM and 6.4 nM for cisplatin, gemcitabine and paclitaxel, respectively (Genomics of Drug Sensitivity in Cancer Database, Wellcome Sanger Institute).

**Figure 2.**
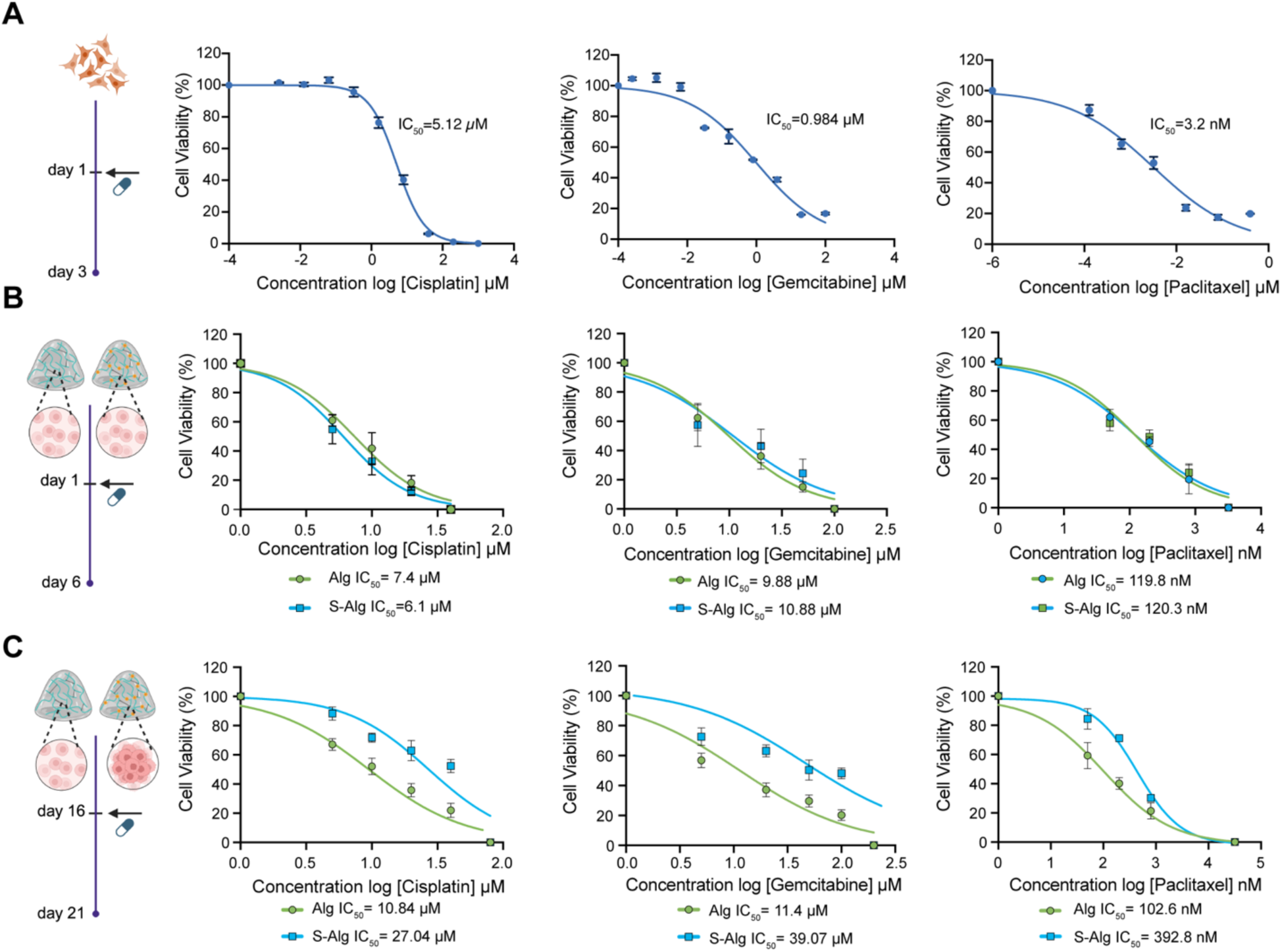
Growth in 3D Sulfated Environment Potently Reduces Tumor Cell Sensitivity to Chemotherapeutics. **A)** Dose-response curves and half-inhibitory concentration (IC50) values for A549 cells after 72h treatment with cisplatin, gemcitabine and paclitaxel in 2D monolayer culture. **B)** Dose-response curves and IC50 values for cisplatin, gemcitabine and paclitaxel for early treatment (day 1) of A549 cells in Alg and S-Alg hydrogels. **C)** Dose-response curves and IC50 values for cisplatin, gemcitabine and paclitaxel for later treatment (day 16) of A549 cells grown in Alg and S-Alg hydrogels. For all drug treatment set-ups, three independent replicates were measured for each drug concentration and results were normalized to the control group. Dose-response curves were assessed with non-linear regression curve fit analysis.

To assess the IC_50_ values of drugs on tumor cells in 3D hydrogels and reveal microenvironment-dependent differences in drug responses, A549 cells were encapsulated in Alg and S-Alg hydrogels and treated with the drugs at indicated doses on day 1 (**Figure 2B**). Early-dose-response screening curves yielded similar profiles and IC_50_ values for all drugs in Alg and S-Alg hydrogels, which demonstrated that modification of alginate did not directly alter drug pharmacokinetics. Furthermore, cisplatin release from Alg and S-Alg hydrogels was assessed with quantification of platinum using inductively coupled plasma mass spectrometry (ICP-MS) confirming that cisplatin diffusion was not affected from sulfate modification of the polymer (**Supplementary Figure 1**). Interestingly, IC_50_ values for early treatment in 3D were close to 2D for cisplatin whereas for gemcitabine and paclitaxel, a sharp increase in IC_50_ was observed indicating a drug-specific effect of the 3D microenvironment on tumor cell responses (**Figure 2A, B**).

A549 cells were next allowed to grow in both Alg and S-Alg hydrogels for 16 days followed by drug treatment for 5 days to assess drug responses after clump formation and morphological adaptation to different matrices (**Figure 2C**). In contrast with early drug responses, a drastic difference in hydrogels was observed. A549 cells grown in S-Alg matrices exhibited significantly less sensitivity to all given drugs. For cisplatin, the IC_50_ value increased from 10.84 µM in Alg hydrogels to 27.04 µM in S-Alg. Gemcitabine and paclitaxel responses were altered even more potently. IC_50_ of gemcitabine increased from 11.4 µM in Alg hydrogels to 39.07 µM in S-Alg and for paclitaxel, an increase from 102.6 nM in Alg hydrogels to 392.8 nM in S-Alg was observed. Overall, adaptation of A549 cells to sGAG-mimetic tumorigenic environment significantly altered the sensitivity to chemotherapy.

### 2.3. Sulfated Environment Promotes Stress Compensation in Lung Tumor Cells

Followed by establishment of dose-response curves for each drug and condition, A549 cells grown in Alg/S-Alg hydrogels were next treated with a single chosen dose of cisplatin (10 µm), gemcitabine (5 µM) or paclitaxel (50 nM) at early (day 1) or late (day 16) time-points and assessed for metabolic activity. For early time-point treatments, the metabolic activity of A549 cells were quite similar in both Alg and S-Alg hydrogels for all three drugs (**Supplementary Figure 2**). However, when treatments were given at day 16 after allowing cellular growth and adaptation within hydrogels, the metabolic activity levels of treated A549 cells were significantly higher in sulfated hydrogels for all three drugs (**Figure 3A**).

**Figure 3.**
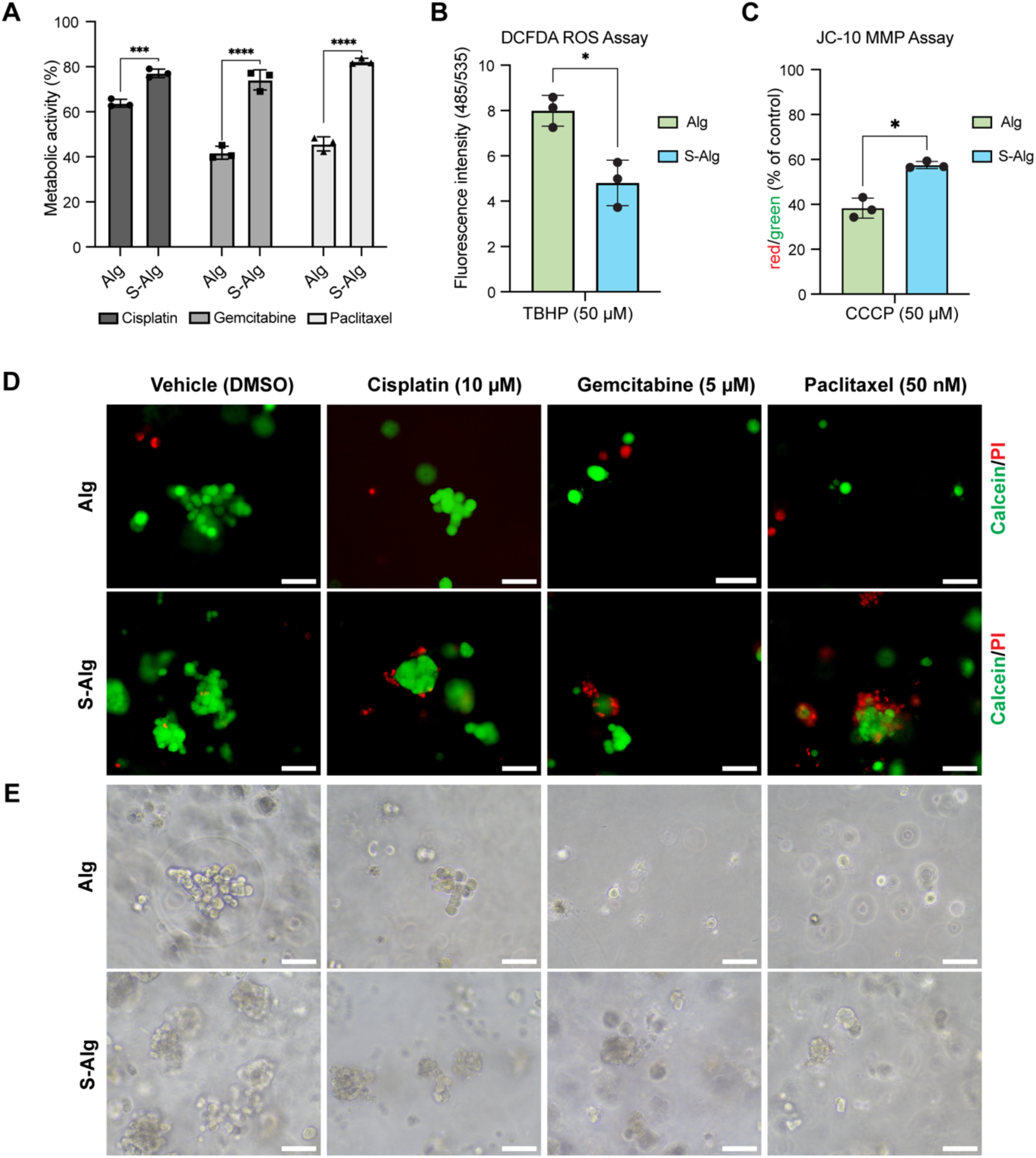
Sulfated Environment Promotes Stress Compensation in Lung Tumor Cells. **A)** Metabolic activity of A549 cells encapsulated in Alg or S-Alg hydrogels for 16 days followed by treatment with cisplatin (10 µM), gemcitabine (5 µM) or paclitaxel (50 nM) for 5 days. Three independent replicates were measured and error bars indicate mean ± SD. Statistical analysis was performed using two-tailed t-test, *p<0.05; **p<0.01; ***p<0.001. **B)** DCFDA ROS production assessment of A549 cells after treatment with 50 µM TBHP for 4 hours in Alg and S-Alg hydrogels. Three independent replicates were measured, and error bars indicate mean ± SD. Statistical analysis was performed using two-tailed t-test, *p<0.005. **C)** JC-10 mitochondrial membrane potential assessment of A549 cells after treatment with 50 µM CCCP in Alg and S-Alg hydrogels. Three independent replicates were measured, and error bars indicate mean ± SD. Statistical analysis was performed using two-tailed t-test, *p<0.05. **D)** Calcein-AM (green) and Propidium iodide (PI) (red) staining of A549 cells encapsulated in Alg or S-Alg hydrogels for 16 days followed by treatment with cisplatin (10 µM), gemcitabine (5 µM) or paclitaxel (50 nM) for 5 days. Scale bar: 70 µm. **E)** Bright-field microscopy images of cells encapsulated in Alg or S-Alg hydrogels for 16 days followed by treatment with cisplatin (10 µM), gemcitabine (5 µM) or paclitaxel (50 nM) for 5 days. Scale bar: 70 µm.

To evaluate microenvironment-induced differences in oxidative stress response in A549 cells, we quantified reactive oxygen species (ROS) levels in Alg and S-Alg hydrogels following treatment with tert-butyl hydroperoxide (TBHP), a well-established inducer of oxidative stress (**Figure 3B**). We employed the fluorogenic probe DCFD, which measures hydroxyl and peroxyl radicals as well as other intracellular ROS. Upon diffusion into cells, DCFDA is deacetylated by cellular esterases to a non-fluorescent form whose oxidation by ROS yields the highly fluorescent DCF. Fluorescence intensity at 485/535 nm was measured to monitor ROS levels in A549 cells in Alg and S-Alg hydrogels. Cells in S-Alg hydrogels demonstrated a significant decrease in TBHP-induced generation of ROS compared to Alg hydrogels indicating increased tolerance for apoptosis-promoting stress factors.

Disruption of mitochondrial membrane potential (MMP) is one of the key early events in the apoptosis pathway [52]. Loss of MMP enables the release of cytochrome c from mitochondria into the cytosol and activates downstream caspase signaling. Many chemotherapeutic agents promote apoptosis through mitochondrial destabilization. Therefore, we performed MMP assessment as a measure of cellular stress and apoptotic susceptibility. To evaluate mitochondrial integrity, we adapted the JC-10 assay to A549 cells encapsulated in Alg and S-Alg hydrogels and measured MMP following treatment with carbonyl cyanide *m*-chlorophenyl hydrazone (CCCP), a commonly used mitochondrial uncoupler (**Figure 3C**). JC-10, a cationic and lipophillic dye, exhibits a shift in its emission spectrum according to the state of MMP. In healthy cells with intact MMP, JC-10 forms red-emitting aggregates on mitochondrial membrane. In apoptotic cells, MMP disruption leads to its diffusion and dissipation into green-emitting monomeric form. A549 cells cultured with S-Alg hydrogels displayed significantly higher red-to-green fluorescence ratio indicating higher mitochondrial integrity under CCCP-induced stress when compared to Alg hydrogels. Together with the diminished ROS levels, these results highlight the capacity of S-Alg hydrogels to provide a stress-compensating microenvironment for lung tumor cells.

Next, we evaluated the viability and spatial organization of A549 cells in Alg and S-Alg hydrogels upon treatment at day 16 with cisplatin (10 µM), gemcitabine (5 µM) or paclitaxel (50 nM) with calcein/PI staining (**Figure 3D**). Cells cultured in S-Alg hydrogels formed clusters characterized by a viable core and more peripheral localization of PI-positive cells. Notably, a central necrosis, typically observed for clumps larger than 100 µm upon lack of oxygen and nutrient diffusion, was not observed for the clumps in S-Alg hydrogels. In contrast, A549 cells in Alg hydrogels organized as single cells or non-cohesive clusters with both calcein- and PI-positive cells interspersed throughout. Bright-field imaging further complemented the data revealing morphological indicators of stress upon drug treatment for cells in Alg hydrogels such as membrane irregularities, whereas cells in S-Alg hydrogels maintained a more stable epithelial-like morphology and cluster structure **(Figure 3E)**.

These findings demonstrate that aberrant sulfation in the extracellular microenvironment mitigates oxidative and mitochondrial stress and supports higher survival under chemotherapeutic challenge.

### 2.4. Increased Sulfation in the ECM Regulates Expression of Genes Involved in Apoptosis

Apoptosis proceeds through two main routes, the extrinsic, death receptor-mediated pathway and the intrisic, mitochondria-dependent pathway [53]. The extrinsic pathway involves ligand binding to death receptors belonging to the tumor necrosis factor (TNF) receptor gene superfamily [54]. Among these, the Fas receptor (FasR) is one of the most extensively characterized death receptors. Binding of the Fas ligand (FasL) induces recruitment of adaptor proteins through its cytoplasmic death domain, thereby activating downstream apoptotic signaling [53]. The intrinsic pathway is activated by mitochondrial outer membrane permeabilization and subsequent release of cytochrome c, a process tightly regulated by the members of the Bcl-2 protein family [55]. Both pathways converge on the activation of executioner caspase-3 through proteolytic cleavage, making cleaved caspase-3 a widely used marker for apoptosis [53]. To determine the effect of aberrant ECM sulfation on apoptotic signaling, we performed immunofluorescence (IF) staining for cleaved caspase-3 on A549 cells in Alg and S-Alg hydrogels in response to treatment with chemotherapeutics (**Figure 4A**). Phalloidin/DAPI staining (gray/blue) was used as counterstaining to visualize the morphology and integrity of multicellular clusters, while red fluorescence indicated cleaved caspase-3 expression. A549 cells encapsulated in Alg hydrogels showed markedly elevated levels of cleaved caspase-3 following exposure to cisplatin, gemcitabine, and paclitaxel (**Figure 4B**). In contrast, cells in S-Alg hydrogels exhibited reduced caspase-3 activation in line with enhanced mitochondrial stability and improved viability observed in **Figure 3**.

**Figure 4.**
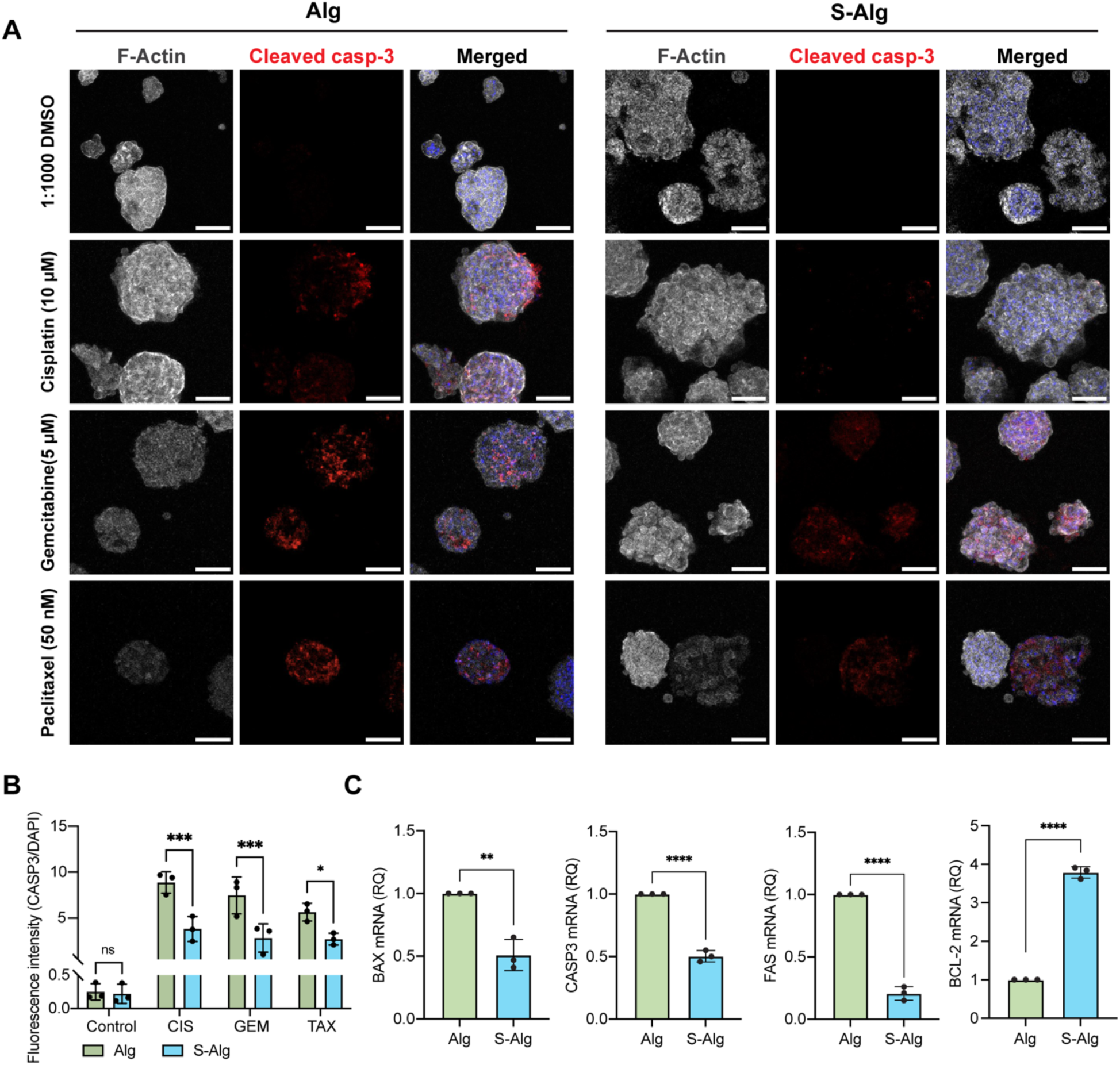
Increased Sulfation in the ECM Suppresses Apoptotic Markers in Lung Tumor Cells. **A)** Representative immunofluorescence staining images for apoptotic marker cleaved caspase-3 expression of A549 cells in Alg and S-Alg hydrogels. Cleaved caspase-3 (red); phalloidin (gray); DAPI (blue). Scale bar: 100 µm. **B)** Quantification of cleaved caspase-3 fluorescence intensity between Alg and S-Alg hydrogels. Data are shown as mean ± SD of three replicates (the average of 10-30 images per replicates). Statistical analysis was performed using two-tailed t-test, ***p<0.001, *p<0.05. **C)** Expression level of genes involved in regulation of apoptosis. Gene expression analysis was performed following 3-week culture of A549 cells in Alg and S-Alg hydrogels. Three independent replicates were measured, and error bars indicate mean ± SD. Statistical analysis was performed using two-tailed t-test, **p<0.01; ****p<0.0001.

Next, we sought to elucidate whether sulfation alters the expression of key apoptotic markers in lung tumor cells (**Figure 4C**). A549 cells in Alg hydrogels demonstrated significantly higher expression of the pro-apoptotic gene *BAX* and the executioner caspase gene *CASP3*. On the other hand, expression of the anti-apoptotic *BCL-2* gene was upregulated for cells in S-Alg hydrogels. Additionally, expression of the death receptor gene *FAS* was observed to be significantly higher for cells in Alg hydrogels, suggesting increased sensitivity for death ligand-mediated apoptosis in this matrix microenvironment.

Together with our cleaved caspase-3 IF data, these alterations in transcriptional profiles show that S-Alg hydrogels suppress both intrinsic and extrinsic apoptotic signaling programs in lung tumor cells and support activation of pro-survival pathways.

### 2.5. Sulfated ECM Acts Through the PI3K/Akt-ABCB1 Axis and Modulates Drug-Specific Response

Given the sulfation-mediated suppression in apoptotic signaling observed in lung tumor cells, we next sought to identify the upstream pathways enabling reduced drug sensitivity in sulfated matrices. In our previous work, we demonstrated that aberrant sulfation in the lung tumor ECM enhances PI3K/Akt activation [43], a pathway established for promoting survival and evasion from apoptosis. As PI3K/Akt signaling has been widely implicated in the acquisition of multidrug resistance (MDR) as well as transcriptional regulation of ATP-binding casette (ABC) efflux transporters, we next evaluated whether this signaling axis contributes to the attenuated chemotherapeutic response observed in S-Alg hydrogels.

Among the major drug efflux pumps associated with chemotherapy resistance, ABCB1 (P-glycoprotein) is known to be directly modulated by PI3K/Akt activation [56]. Consistent with PI3K pathway activation in sulfated matrices [43], A549 cells encapsulated S-Alg hydrogels exhibited significantly higher ABCB1 expression compared to Alg hydrogels (**Figure 5A, B**). To examine whether sulfation broadly affected the expression of ATP-dependent drug efflux pumps, we analyzed additional transporters which revealed either nonsignificant change or decreased gene expression in S-Alg hydrogels (**Supplementary Figure 3**), suggesting that ABCB1 is the dominant efflux pump upregulated in the sulfated microenvironment.

**Figure 5.**
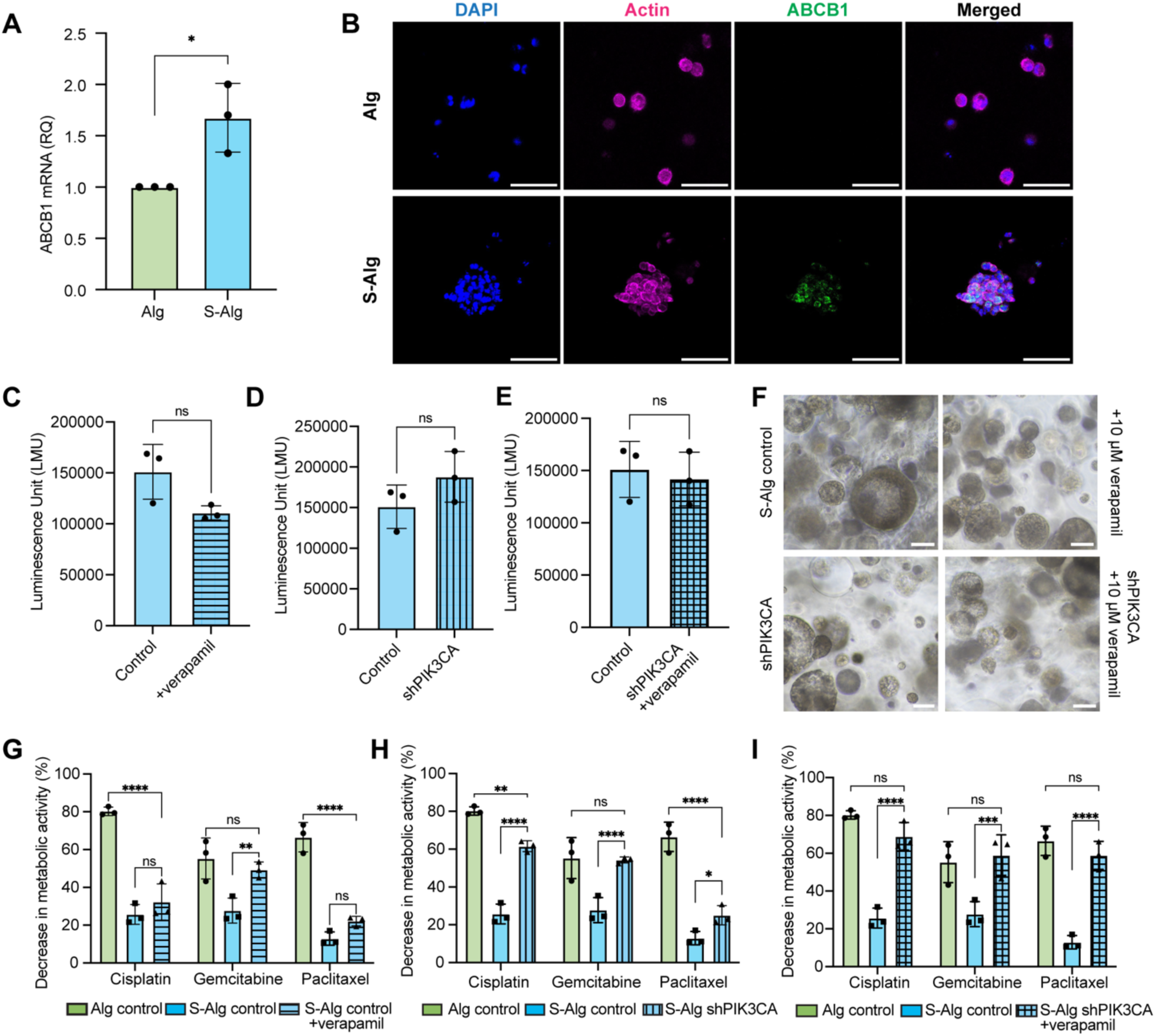
Sulfated ECM Acts Through the PI3K/Akt-ABCB1 Axis and Modulates Drug-Specific Response. **A)** Expression levels of *ABCB1* gene in A549 cells grown in Alg and S-Alg hydrogels. Three independent replicates were measured, and error bars indicate mean ± SD. Statistical analysis was performed using two-tailed t-test, *p<0.05 **B)** Representative immunofluorescence staining images for ABCB1 in Alg and S-Alg hydrogels. ABCB1 (green); Actin (magenta); DAPI (blue). Scale bar: 100 µm. Metabolic activity assessment of A549 cells in S-Alg hydrogels under **C)** verapamil treatment, **D)** *PIK3CA* knockdown, **E)** concurrent verapamil treatment and *PIK3CA*-knockdown conditions. Three independent biological replicates were measured, and error bars indicate mean ± SD. Statistical analysis was performed using two-tailed t-test, ns not significant. **F)** Bright-field microscopy images for A549 cells grown in S-Alg hydrogels under untreated, verapamil-treated, PI3K knockdown and dual inhibition conditions. Scale bar=50 µm. Metabolic activity assessment of A549 cells in S-Alg hydrogels after treatment with cisplatin (10 µM), gemcitabine (5 µM) and paclitaxel (50 nM) under **G)** verapamil treatment, **H)** *PIK3CA*-knockdown, **i)** concurrent verapamil treatment and PIK3CA knockdown conditions. Three independent replicates were measured; and statistical analysis was performed by using One-way ANOVA, ns not significant; *p<0.05; **p<0.01; ***p<0.001; ****p<0.0001.

Next, functional inhibition of ABCB1 and PI3K was pursued to determine their involvement in sulfation-induced decrease in drug sensitivity. ABCB1 activity was inhibited using verapamil and metabolic activity of A549 cells in S-Alg hydrogels was measured following treatment (**Figure 5C**). In parallel, shRNA-mediated knockdown of *PIK3CA* in A549 cells was performed to test the effect of reduced PI3K activity in S-Alg hydrogels (**Figure 5D**). The combined effect of PI3K downregulation and ABCB1 inhibition was assessed by treating *PIK3CA*-knockdown cells with verapamil (**Figure 5E**). Metabolic activity of A549 cells in S-Alg hydrogels across all conditions, verapamil-treatment, *PIK3CA*-knockdown and combination, did not differ significantly from untreated controls in the absence of chemotherapeutics (**Figure 5C-E**). Bright-field microscopy images confirmed that all groups demonstrated comparable growth and morphology prior to drug exposure (**Figure 5F**), enabling controlled, mechanism-focused further assessment of chemotherapeutic responses.

We then evaluated whether ABCB1 inhibition of PIK3CA knockdown sensitizes A549 cells in S-Alg hydrogels to cisplatin, gemcitabine and paclitaxel. A549 cells in S-Alg hydrogels remained largely unresponsive to cisplatin and paclitaxel when ABCB1 was inhibited individually (**Figure 5G**). Interestingly, sensitivity to gemcitabine increased to levels seen in Alg hydrogels when either ABCB1 or PI3K was inhibited alone, suggesting that nucleoside analog-based therapies are more acutely impacted by ABCB1 modulation (**Figure 5G, H**). Inhibition of PI3K alone strongly increased cisplatin sensitivity in S-Alg hydrogels which was still significantly lower compared to Alg hydrogels (**Figure 5H**). On the other hand, PI3K inhibition alone had a much weaker effect on paclitaxel response and caused a small increase in sensitivity in S-Alg hydrogels (**Figure 5H**). Moreover, dual inhibition of PI3K and ABCB1 completely sensitized A549 cells in S-Alg hydrogels for all three drugs to the levels observed in Alg hydrogels (**Figure 5I**). These findings reveal that the PI3K/Akt-ABCB1 axis contributes synergistically to the impaired chemotherapeutic response in sulfated matrices and that dual targeting is required for reversing this phenotype for certain drug classes which can be attributed to their distinct mechanism of action. Our results support a model in which sGAG-mimetic cues promote survival and strengthening of drug efflux capacity through PI3K activation leading to drug-specific modulation of chemoresistance.

## 3. Discussion

The advent of 3D culture systems has reshaped our understanding of how TME regulates tumor development, progression and therapeutic response. By incorporating biophysical and biochemical microenvironmental parameters absent in 2D models, engineered 3D matrices enable investigation of tumor-specific changes in ECM cues that drive malignant phenotypes [25]. In our recent work, we demonstrated that aberrant sGAG deposition in the lung TME stimulates cancer stemness and invasiveness in lung adenocarcinoma [43], motivating a deeper investigation of how ECM sulfation modulates therapeutic vulnerability.

Here, we show that a tumor-mimetic hydrogel engineered to represent the elevated sGAG levels of lung tumor matrices activates compensatory survival programs, supports stress tolerance and regulates response to chemotherapeutics in lung adenocarcinoma cells. We employed sulfated alginate matrices with stiffness tuned to match unmodified alginate hydrogels and isolated sulfation as a single biochemical variable. Within this controlled system, tumor-mimetic S-Alg hydrogels enhanced proliferation and survival, suppressed early apoptotic events, stabilized mitochondrial function, reduced oxidative stress and decreased sensitivity to cisplatin, gemcitabine and paclitaxel. These findings align with our previous work linking sGAG-rich matrices to stem-like phenotype [43], and further establish ECM sulfation as an important determinant of drug response in lung adenocarcinoma.

A key advance of this study is the demonstration that sulfation alone, independent of aberrant ECM mechanics, regulates response to chemotherapeutic agents with different mechanisms of action. Cisplatin (alkylating agent), gemcitabine (anti-metabolite) and paclitaxel (microtubule stabilizer) possess well-characterized, distinct mechanisms for resistance in tumor cells [57]. Intriguingly, all three drugs showed significantly attenuated efficacy in sulfated hydrogels which drove us to resolve the upstream pathways that link sulfation to drug-specific resistance phenotypes.

Sulfation suppresses ROS accumulation upon THBP challenge and preserves MMP under CCCP stress, two upstream checkpoints for intrinsic apoptosis. Cleaved caspase-3 expression is downregulated in S-Alg hydrogels compared to Alg hydrogels in line with a transcriptional shift towards an anti-apoptotic profile. These features are consistent with maintenance of cohesive multicellular architecture and absence of core necrosis which emphasize both biochemical (oxidative/mitochondrial) and biophysical (cluster integrity) means of stress buffering in the sulfated microenvironment. Notably, early dose-responses as well as drug diffusion rates are comparable between Alg and S-Alg hydrogels indicating that the late-emerging decrease in drug sensitivity is not a result of altered drug availability but microenvironmental adaption.

Sulfated hydrogels induced a selective upregulation of ABCB1, a known efflux protein associated with paclitaxel and gemcitabine resistance [57, 58]. Conversely, expression of cisplatin-related efflux transporters such as ABCC1 and ATP7B did not increase with matrix sulfation. Inhibition of ABCB1 activity in S-Alg hydrogels rescued gemcitabine response to levels observed in Alg hydrogels in line with its known susceptibility to efflux modulation as opposed to cisplatin and paclitaxel which did not show an effect. PI3K/Akt signaling acts as an integrator of ECM-mediated survival, chemoresistance and transcriptional regulator of efflux proteins including ABCB1 [56–59]. shRNA-mediated knockdown of PI3K signaling improved responses to all chemotherapeutics in sulfated hydrogels with varying effects among the drugs. Gemcitabine sensitivity returned to Alg levels, cisplatin response improved substantially whereas paclitaxel response increased only modestly. Dual inhibition of both PI3K and ABCB1 fully sensitized cells in S-Alg hydrogels to all three drugs to control levels suggesting that sulfation promotes a signaling-efflux axis which must be targeted simultaneously to reverse certain resistance phenotypes. Limited rescue of paclitaxel response with PI3K or ABCB1 inhibition alone suggests a compensatory mechanism. PGs with their sulfated domains are known modulators of microtubule stability and cytoskeletal organization [40]. This raises the possibility of the contribution of sulfation-dependent cytoskeletal regulation to paclitaxel resistance partially independent of PI3K signaling or efflux alone. The requirement for combined pathway suppression for paclitaxel sensitivity emphasizes the importance of additional PG-mediated regulatory nodes relevant for microtubule-targeting therapies.

Our findings support a growing body of literature showing that the TME can drive malignant transformation and therapy response. Seminal studies demonstrated that ECM mechanics and composition can influence malignancy and invasion [29]. Recent work by LeSavage et al. revealed that increased matrix stiffness promotes chemoresistance in pancreatic cancer and proposed a potential combination of mechanotherapeutics with anti-cancer agents [32]. Similarly, ECM remodeling via lysyl oxidases (LOX) has been shown to promote chemoresistance in triple negative breast cancer (TNBC) and inhibition of LOX or its downstream mediators re-sensitized tumors to treatment [60]. These studies collectively point to the role that ECM-provided cues have in regulation of survival, plasticity, invasion and drug response.

PGs are essential to the aforementioned processes. Their sGAG chains control growth factor availability, RTK activation and downstream signaling including PI3K/Akt [42, 43, 45]. Aberrant expression of PGs is common across cancers [40] and multiple therapeutic strategies targeting PGs or their synthesis have entered preclinical testing [61–63]. These approaches include PG-targeted CAR-T cells, PG-primed dendritic cells, antibody-drug conjugates and inhibitory peptides [41, 64, 65]. Manipulation of heparan sulfate proteoglycans (HSPGs) on ECM through cleavage of their sGAG side chains via heparanases has been explored as a treatment strategy [66]. Dieter et al. demonstrated suppression of heparan sulfation to overcome resistance to MAPK inhibitors in melanoma cells [48]. Although dysregulation of PG metabolism has been shown to correlate with tumor growth and metastasis, their role in tumor progression remains context-dependent due to their structural heterogeneity [44] underscoring the importance of dissecting their specific features such as sulfation degree.

In summary, sulfated alginate hydrogels provided a chemically defined system to recapitulate the PG abundance of lung tumors and demonstrated that extracellular sulfation actively regulates chemotherapeutic response in lung adenocarcinoma. Our work identifies sulfation as a driver of stress tolerance, apoptosis suppression and decreased drug response mediated by the PI3K/Akt-ABCB1 axis in a drug-specific manner. Our findings position PGs/sGAGs as compelling therapeutic targets and support future strategies for personalized, combination-based approaches for sensitization of lung adenocarcinoma patients to chemotherapeutics.

## 4. Conclusion

In this work, we establish extracellular matrix sulfation as a potent regulator of chemotherapeutic response in lung adenocarcinoma. By isolating sulfation within a stiffness-controlled and inert 3D hydrogel system, we demonstrate that elevated sGAG cues in tumor matrices are sufficient to promote survival, cellular stress tolerance, suppression of apoptosis and drug sensitivity through PI3K/Akt signaling and ABCB1-mediated efflux. Distinct patterns of sensitization across cisplatin, gemcitabine, and paclitaxel emphasize that matrix sulfation modulates therapy response in a drug-specific manner. These findings highlight aberrant PG/sGAG deposition in tumors as active participants of resistance development and targetable determinants of treatment efficacy which support development of personalized and ECM-informed strategies to improve patient outcomes in lung cancer.

## 5. Experimental Section Modification of alginate

Sulfation of alginate (Novamatrix, Norway) was carried out as previously described [52]. Briefly, 99% chlorosulfonic acid (HSO_3_Cl; Sigma) was diluted in formamide (Sigma) at 2% (v/v) and added dropwise onto alginate. The reaction was carried out at 60 °C with agitation for 2.5 h. Sulfated alginate was precipitated with cold acetone, and dissolved in deionized water and adjusted to neutral pH. The solution was then dialyzed with a 12000 Da molecular weight cut-off (MWCO) tube (Sigma) in 100 mM NaCl for 48 h and then deionized water for 72 h and lyophilized.

### Chemical Characterization of Modified Alginate

Elemental analysis of sulfur content was done by high-resolution inductively coupled mass spectrometry (ICP-MS) analysis on alginate dissolved in 0.1 M HNO_3_. Degree of sulfation (DS), number of sulfate groups per monomer, was calculated using the mass balance equation assuming one sodium counterion for each negatively charged group and one water molecule per monosaccharide: monosaccharide mass = C_6_O_6_H_5_ + (DS + 1) Na^+^ + (DS)SO ^-^ + H_2_O. Sulfated alginate with a DS value of 0.35 was used in the study.

### Mechanical Characterization

Mechanical characteristics of hydrogels were assessed with a rheometer (Malvern Kinexus) with parallel plate geometry (diameter: 10 mm). Hydrogel precursor was poured onto the lower plate and the upper layer was lowered with a measuring gap adjusted to 1 mm. Mineral oil (Sigma) was used at the edges of the gel to prevent dehydration during measurement. Oscillatory rheology was used to monitor hydrogel storage modulus at constant frequency and amplitude (1 Hz; 1% strain) for 2 h. All measurements were done in triplicates.

### Cell Culture

Human lung adenocarcinoma cell line A549 (CCL-185) was purchased from American Tissue Culture Collection (ATCC) and cultured in growth medium DMEM/F12 (Lonza) supplemented with 10% fetal bovine serum (FBS) (Biowest) and 1% penicillin-streptomycin (P/S) (Gibco) with incubation at 37 °C and 5% CO_2_. shRNA-mediated knockdown of *PIK3CA* in A549 cells was carried out as previously described [43].

### Cell Encapsulation in Hydrogels

Prior to encapsulation, cells were trypsinized, washed, resuspended and counted. Alginate and alginate sulfate solutions were prepared in serum free DMEM/F12 medium and sterile filtered (0.2 µm pore size). Cell suspension (2 x 10^5^ cells/ml final density), alginate/alginate sulfate and CaSO_4_ solutions were speedily combined using double Luer-lock syringes and casted onto wells of a 24-well plate. The samples were incubated at 37 °C for 45 min for complete gelation after which complete growth medium was added. The hydrogels were cultured up to 21 days with medium change every 3 days.

### Quantification of dsDNA

DNA content of cells encapsulated within alginate/alginate sulfate hydrogels was determined to confirm cell growth. Gels were washed three times with wash buffer (150 x 10^-3^ M NaCl and 5 x 10^-3^ M CaCl_2_) before collection for assay. Triplicates of gels were collected for each time points (day 0, 7, 14, 21). Gels were stored at -80 °C until all samples were collected. Then, the hydrogels were digested with papain (125 µg/ml, Sigma) in 10 x 10^-3^ M EDTA, sodium phosphate (100 x 10^-3^ M, Sigma) and L-cysteine (10 x 10^-3^ M, Sigma) at pH 6.3 and incubated at 60 °C overnight and vortexed several times. Quant-iT PicoGreen (Invitrogen) dsDNA assay was used to quantify dsDNA according to the manufacturer’s protocol. Briefly, diluted samples were mixed with Quant-iT PicoGreen reagent after digestion and incubated for 5 min at room temperature. Fluorescence was measured at 520 nm with excitation at 485 nm using a microplate reader (BioTek Synergy).

### Assessment of Cell Viability and Morphology

Cell viability was assessed with Calcein/AM (Invitrogen) and propidium iodide (Sigma) staining. Briefly, A549 cells encapsulated in hydrogels were stained with 2 x 10^-6^ M Calcein-AM and 30 µg/ml propidium iodide in growth medium for 1 h at 37 °C in dark conditions. Then, gels were washed twice with a growth medium and imaged with a fluorescent microscope (Nikon ECLIPSE Ts2). Cell morphology was assessed with F-actin and DAPI staining. Hydrogels were collected at given time points, fixed with 4% paraformaldehyde for 45 min at room temperature and washed three times with 5 min intervals. Then, permeabilization and blocking were achieved with incubation in BSA (5%, Sigma) and Triton-X-100 (1%, Sigma) for 1 h at room temperature. For F-actin staining, the hydrogels were incubated with Phalloidin-iFluor 488 Reagent (1:1000, Abcam) for 45 min at room temperature followed by three 5 min-washes. Nuclei counter staining was performed with DAPI (1µg/ml, Sigma). Confocal microscopy (Leica SP8) was used for imaging and at least three hydrogels were used per condition.

### Metabolic Activity Assay

CellTiter-Glo 3D^®^ Cell Viability Assay (Promega) was performed in hydrogels according to manufacturer’s instructions with minor modifications. Hydrogels were incubated with assay buffer in a 48-well plate for 45 min at room temperature following a 10 min agitation with a plate shaker. Luminescence was measured for at least three hydrogels using a microplate reader (BioTek Synergy).

### Drug Response Assessment

For measurement of 2D drug response, CellTiter-Glo Luminescent Cell Viability Assay (Promega) was performed according to the manufacturer’s instructions. Cells were seeded onto 96-well black wall plates at a seeding density of 2,000 cells per well. After 24h, cells were treated with cisplatin (232120, Sigma), gemcitabine (Y0000675, European Pharmacopoeia) and paclitaxel (Y0000698, European Pharmacopoeia). Half inhibitory concentrations (IC_50_) of drugs were determined after 72h treatment on a range of cisplatin doses (1 mM to 2.56 nM), gemcitabine doses (100 µM to 0.256 nM), and paclitaxel doses (400 nM to 0.128 nM) with 1:5 serial dilutions.

For measurement of 3D early and late drug responses, cells were encapsulated in hydrogels with 5x10^5^ cells/ml and 2 x 10^5^ cells/ml seeding densities, respectively. In early response, cells were treated with drugs after 24h of culture. For the late response, cells were cultured in hydrogels for 16 days followed by drugs administration for 5 days. IC_50_ values were determined on a range of cisplatin doses (300 µM to 2.4 µM), gemcitabine doses (200 µM to 5 µM) and paclitaxel doses (3.2 µM to 50 nM) with CellTiter-Glo 3D^®^ Cell Viability Assay (Promega) as described above. DMSO without drugs was used in non-treatment control for gemcitabine and paclitaxel to normalize the cell viability. Cisplatin was solved in saline solution, and its control group was prepared accordingly. The results were analyzed on GraphPad Prism 10 and IC_50_ values were assessed using a nonlinear regression curve fit.

### DCFDA ROS Assay

For measurement of reactive oxygen species in cells, DCFDA/ROS assay (Abcam) was used. A549 cells were encapsulated in Alg and S-Alg hydrogels and cultured for 21 days. Then, 50 µM TBHP treatment was performed for 4 hours to trigger the release of ROS. At the end of treatment, cells were retrieved from hydrogels and seeded onto a clear-bottom black-side 96-well plate with a density of 1.5x10^5^ cells/well. After that point, the manufacturer’s instructions were followed. Fluorescence intensity was measured at 535 nm with an excitation at 485 nm using a microplate reader (BioTek Synergy).

### JC-10 Mitochondrial Membrane Potential (MMP) Assay

JC-10 MMP Assay Kit (Abcam) was used to measure the MMP. A549 cells were encapsulated in Alg and S-Alg hydrogels and cultured for 21 days. Then, 50 µM CCCP (Sigma) treatment was performed for 4 hours. At the end of treatment, cells were retrieved from hydrogels and seeded onto a clear-bottom black-side 96-well plate with a density of 10x10^3^ cells/well. After that point, the manufacturer’s instructions were followed. Fluorescent intensities were measured at 525 nm with an excitation at 490 nm and 590 nm with an excitation at 540 nm. Ratio between red/green fluorescence intensity was determined as a measure of MMP.

### Real Time-Quantitative Polymerase Chain Reaction (qRT-PCR)

TRIzol Reagent (Invitrogen) was added onto the hydrogels followed by crushing with a pestle on dry ice. Then, the samples were centrifuged at 12000 rpm for 5 min at 4 °C, chloroform was added to the supernatant and incubated on ice for 10 min followed by centrifugation at 12000 rpm for 15 min at 4 °C. Aqueous phase was collected onto which 70% EtOH was added in equal volume. RNA isolation was performed with Nucleospin RNA II (MN) kit following manufacturer’s instructions. Reverse transcription was performed with M-MLV Reverse Transcriptase Kit (Invitrogen) using 1 µg RNA. qPCR was performed on a LightCycler equipment (Roche) using SYBR Green (Roche) dye. Primer sequences that were used are listed in **Table S1, Supporting Information**.

### Immunofluorescence Staining

Hydrogels were fixed with 4% PFA for 45 min at room temperature. Next, the gels were washed three times for 5 min with wash buffer followed by permeabilization and blocking in 5% goat serum (Gibco), 1% BSA, 1% Triton X-100 for 1 h at room temperature. Incubation with a primary antibody against cleaved caspase-3 (1:500, Cell Signaling Technologies, 9664S) was performed overnight at 4°C in staining buffer containing 1% BSA and 0.1% Triton X-100. Following primary antibody incubation, the hydrogels were washed twice for 10 min at room temperature with staining buffer and then incubated with the secondary antibody overnight at 4°C using a goat anti-rabbit Alexa Fluor 555-tagged (1:200, Abcam) antibody. The hydrogels were washed at room temperature for 2h, treated with DAPI (1 µg/mL) for nuclei counter staining and washed again for 15 min. Confocal microscopy (Leica SP8) was used for the imaging. Immunofluorescence staining for ABCB1 was performed with a primary antibody against ABCB1 (1:200, Cell Signaling Technologies, 13342S) following the same protocol described above.

## Supporting Information

Supporting Information is available.

## Acknowledgements

The authors acknowledge funding from the International Fellowship for Outstanding Researchers Program of TÜBİTAK (118C238) and European Union’s Horizon 2020 research and innovation program under the Marie Skłodowska-Curie grant agreement (101032602). The entire responsibility of the publication belongs to the owner, the financial support from TÜBİTAK does not mean that the content of the publication is approved in a scientific sense by TÜBİTAK. BioRender.com was used to create the icons used in Figures 1 and 2. The authors gratefully acknowledge the use of services and facilities of Koç University Research Center for Translational Medicine (KUTTAM). The authors gratefully acknowledge Dr. Øystein Arlov and Marianne Kjos from Sintef for ICP-MS analyses. The authors acknowledge the use of Koç University Surface Science and Technology Center (KUYTAM) infrastructure for mechanical assessments of hydrogels.

## Author Contributions

E.Ö. conceived the project and obtained funding. E.Ö. and S.S. designed the research. S.S. conducted experiments and analyzed data. S.S. and E.Ö. wrote the manuscript.

## Data Availability Statement

The data supporting the findings of the study are available from the corresponding author upon reasonable request.

