## Supplementary Information for "Engineered Matrices Reveal Sulfation-Mediated Stress Adaptation and Drug-Specific Modulation of Chemotherapeutic Response"

### Extracellular Matrix Sulfation Reprograms Stress Tolerance and Drug-Specific Chemotherapeutic Response

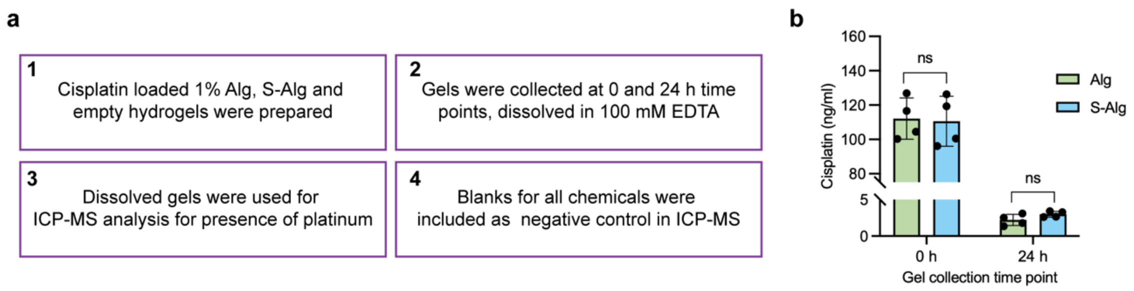


**Supplementary Figure 1.** Inductively coupled plasma mass spectrometry (ICP-MS) analysis for cisplatin release in Alg and S-Alg. **a)** Sample collection steps for ICP-MS analysis. **b)** Quantification of the cisplatin concentration (ng/ml) based on platinum (Pt) elemental analysis in hydrogels at 0 hour (after crosslinking) and 24 hours after aspiration of culture medium. Three independent replicates were measured, and error bars indicate mean ± SD. Statistical analysis was performed using two-tailed t-test, ns not significant.

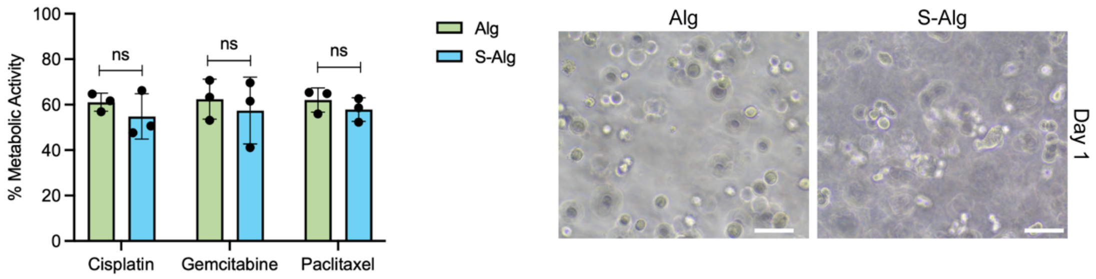


**Supplementary Figure 2.** Early drug response of A549 cells in Alg and S-Alg hydrogels for cisplatin (10 μM), gemcitabine (5 μM) and paclitaxel (50 nM) treatments. Cells were encapsulated with 5x10$^{5}$ cells/ml seeding density and drugs were given one day after cell encapsulation. Cells were treated for 5 days, and metabolic activity was measured. Three independent replicates were measured, and error bars indicate mean ± SD. Statistical analysis was performed using two-tailed t-test, ns not significant. BF images show the appearance of gels at treatment day, scale bar: 70 μm.

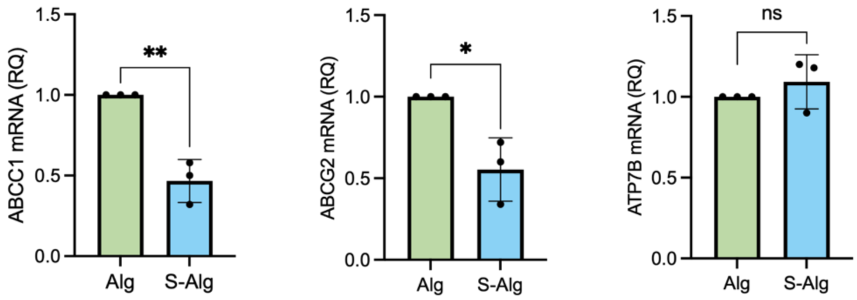


**Supplementary Figure 3.** Expression levels of ATP binding cassette (ABC) drug efflux transporters that are involved in development of chemoresistance. Three independent replicates were measured, and error bars indicate mean ± SD. Statistical analysis was performed using two-tailed t-test, ns not significant; *p<0.05; **p<0.01.

**Supplementary Table 1:** qPCR forward and reverse primer sequences used in this study.

| Gene | Forward Primer (Sense) | Reverse Primer (anti-sense) |
|---|---|---|
| ABCB1 | CAGCAAAGGAGGCCAACATA | CTGTCTAACAAGGGCACGAG |
| ABCG2 | GGGTTTGGAACTGTGGGTAG | GTGACCTCCCAGAGCTAGAA |
| ABCC1 | ACTGGAATGTCACGTGGAAT | CTGAATGTAGCCTCGGTCAT |
| ATP7B | GGAAACTGCAAGGAGTAGTG | CTTCGGGCTGAATGAGATAAG |
| BCL-2 | GCCAGGGTCAGAGTTAAATAG | CCTCTCTTGCGGAGTATTTG |
| BAX | CTGGACAGTAACATGGAGCT | GCAAAGTAGAAAAGGGCGAC |
| CASP3 | GTGGAGGCCGACTTCTTGTA | ACCCGGGTAAGAATGTGCAT |
| FAS | GTGACCCTTGCACCAAATGT | GAAGACAAAGCCACCCCAAG |
